# True causal effect size heterogeneity is not required to explain trans-ethnic differences in GWAS signals

**DOI:** 10.1101/085092

**Authors:** Daniela Zanetti, Michael E. Weale

## Abstract

Through genome-wide association studies (GWASs), researchers have identified hundreds of genetic variants associated with particular complex traits. Previous studies have compared the pattern of association signals across different populations in real data, and these have detected differences in the strength and sometimes even the direction of GWAS signals. These differences could be due to a combination of (1) lack of power (insufficient sample sizes); (2) minor allele frequency (MAF) differences (again affecting power); (3) linkage disequilibrium (LD) differences (affecting power to ‘tag’ the causal variant); and (4) true differences in causal variant effect sizes (defined by relative risks).

In the present work, we sought to assess whether the first three of these reasons are sufficient on their own to explain the observed incidence of trans-ethnic differences in replications of GWAS signals, or whether the fourth reason is also required. We simulated case-control data of European, Asian and African ancestry, drawing on observed MAF and LD patterns seen in the 1000-Genomes reference dataset and assuming the true causal relative risks were the same in all three populations.

We found that a combination of Euro-centric SNP selection and between-population differences in LD, accentuated by the lower SNP density typical of older GWAS panels, was sufficient to explain the rate of trans-ethnic differences previously reported, without the need to assume between-population differences in true causal SNP effect size. This suggests a cross-population consistency that has implications for our understanding of the interplay between genetics and environment in the aetiology of complex human diseases.

## Introduction

In the past decade, through collaborative efforts and with the aid of genome-wide association studies (GWASs), the genetic basis of common complex diseases, such as Alzheimer’s disease (Medway and Morgan 2014) or type 2 diabetes (Hara et al. 2014), and other complex traits such as height (Wood et al. 2014) or obesity (Stijnen et al. 2014), have been greatly clarified. Most GWAS studies have involved individuals of European origin (96% of all subjects according to one review (Bustamante et al. 2011)), but this picture is changing with increasing efforts to conduct GWASs in Asia (especially China) and Africa. Such studies are important because they allow us to redress the Euro-centric bias in GWASs, and to assess whether signals found in European populations are replicable in other human populations. Trans-ethnic replicability of GWAS signals would imply a common aetiology of complex diseases, and this would have important clinical implications (Visscher et al. 2012) (Marigorta and Navarro 2013). The portability of GWAS results would also allow for ‘trans-ethnic mapping’ to help in the post-GWAS localization of the causal locus, by taking advantage of between-population differences in linkage disequilibrium (LD) (Rosenberg et al. 2010). Conversely, a lack of trans-ethnic replicability, if shown to be linked to true differences in causal risk allele effect sizes, would have implications for the use of genetics in complex disease medicine, both in terms of the portability of polygenic prediction algorithms among populations and in terms of the biological interpretation of GWAS signals, as some degree of population-specific genetic disease aetiology would be implied.

A number of studies have compared genetic association signals across different continental populations. These have found some examples where the observed genetic effects show consistency across ancestral groups, either in terms of effect direction or in the magnitude of the effect (Marigorta and Navarro 2013) (Tan et al. 2010) (Kantor et al. 2013) (N’Diaye et al. 2011), and other examples where GWAS-derived signals point to a pattern of population-specific risk effects (Xue et al. 2013) (Wojczynski et al. 2013), or to a different magnitude in the association signal across populations (Waters et al. 2010) (Thomas et al. 2012). In one such study, Ntzani and colleagues (Ntzani et al. 2012) used the catalog of published GWASs previously maintained by the National Human Genome Research Institute (NHGRI), and currently maintained by the European Bioinformatics Institute (EBI), to characterize the frequency and magnitude of between-population differences seen in replication studies of GWAS signals (effect sizes from replication studies were used to minimise the ‘Winner’s Curse’ bias in discovery GWAS signals). A total of 97 associations were evaluated in both European and Asian populations, 24 in both European and African populations, and 13 in all three groups. They found widespread differences in the frequency of risk alleles between Europeans, Asians and Africans, with absolute differences >10% in 75-89% of the three pairwise comparisons, and they also found differences in reported risk allele effect sizes. Indeed, point estimates of effect size were opposite in direction in 18%, 21%, and 38% in the European-Asian, European-African, and Asian-African comparisons, respectively.

The trans-ethnic differences in genetic risk effects catalogued by Ntzani and colleagues (Ntzani et al. 2012) could be due to one or a combination of the following explanations:

1. Between-population genetic architectures (for both risk allele *frequency* and *effect size*) are the same, but differences in power (say due to low sample sizes) prevent detection in other populations;
2. The genetic *effect size* architectures are the same, but differences in risk allele frequency prevent detection in other populations;
3. The genetic *effect size* architectures are the same, but differences in LD patterns prevent detection in other populations (in cases where the ‘lead’ or most significantly associated SNP is not the truly causal SNP);
4. The genetic *effect size* architectures are different – in other words the causal SNP has a different true genetic effect size in different populations.

Note that this categorization requires one to distinguish effect size differences from allele frequency differences, and this requires one to define a frequency-independent measure of effect size. Here, we use genotypic relative risk as our measure of frequency-independent effect size, and we will restrict our simulations to diseases of low prevalence (1% in all simulations) in order to take advantage if the approximate equivalence between relative risk and odds ratios in this scenario.

Biologically, the most interesting explanation is point (4) above, as this implies there are truly different genetic risk architectures among populations. But since points (1)-(3) can also lead to apparent differences in GWAS signals between populations, it is typically not possible to distinguish between the four possible explanations based on real replications of GWASs (as in the study by Ntzani *et al.* (2012)).

Therefore, in the present work, we employed simulations to assess the relative importance of the four explanations listed above for generating trans-ethnic differences in GWAS signals. We generated simulated genomic data in case-control samples of European, Asian and Sub-Saharan African origin. Random loci (SNPs) in the genome were selected to be ‘truly causal’, and we applied equal disease model parameters across the three population groups. Logistic regression analyses and statistical comparisons across the different continental populations were then performed to evaluate the level of consistency across groups, potential differences in allele frequency, and consequently the degree to which points (1)-(3) above, in the absence of point (4), could explain observed trans-ethnic differences in GWAS signals.

## Materials and methods

In order to mimic a replication study following a GWAS (and thus mimic the effect sizes examined by Ntzani *et al.* (2012), phased haplotypes from the 1000 Genomes Phase 1 dataset (Abecasis et al. 2012) were used as input to simulate case-control genotype data. Utah residents (from the CEPH collection) with Northern and Western European ancestry (CEU, n=85), Finns from Finland (FIN, n=93), British samples from England and Scotland (GBR, n=89), Iberians from Spain (IBS, n=14), and Tuscans from Italy (TSI, n=98) were used as input to simulate European samples (EUR, n=379). Han Chinese from Beijing, China (CHB, n=97), Han Chinese South (CHS, n=100), and Japanese from Tokyo, Japan (JPT, n=89) were used to simulate Asian samples (ASN, n=286). Finally, Yorubans from Ibadan, Nigeria (YRI, n=88) were used to simulate West African samples (only one population, YRI, was used for an African grouping, in recognition of the high degree of genetic heterogeneity among different African samples, making a pan-African “AFR” grouping of little relevance in simulating a typical GWAS). Simulated datasets were generated from SNPs with a ‘global’ MAF >1% (as calculated using all samples in the 1000 Genomes Phase 1 dataset). VCFtools (Danecek et al. 2011) and Beagle (Browning and Browning 2007) software were used to extract and transform 1000 Genomes phased data into a format suitable for GWAsimulator.

Simulations were performed via the GWAsimulator software version 2.1 (Li and Li 2008). This program implements a moving-window algorithm to simulate case-control genotype data based on a set of phased input data. It works outwards from the nominated disease locus to generate the case and control datasets, with patterns of LD similar to the input data. A window size of 5 was used for our simulations, meaning that a haplotype of 4 SNPs was used to propose the allele of the next adjacent SNP.

To mimic a Euro-centric bias in the GWAS study forming the basis for this simulated replication study, a ‘true’ causal disease locus was selected at random for each simulation run from the set of all autosomal SNPs in the 1000 Genomes Project Phase 1 dataset with a minor allele frequency (MAF) >5% specifically in the EUR grouping. The Genotypic Relative Risk for this locus was set at 1.3, with a multiplicative effect, setting the alternative (non-reference) allele as the risk allele. Genomic regions of 500 kb were simulated, 250 kb upstream and downstream of the randomly selected disease locus. For each simulation run, 2000 cases and 2000 controls were created with a disease prevalence of 1%. Logistic regression analyses were performed on the simulated data from the three continental groups (EUR, ASN and YRI), using Plink software version 1.07 (Purcell et al. 2007).

Following Ntzani and colleagues (Ntzani et al. 2012), the following metrics were used to assess apparent trans-ethnic differences in our simulated replications of GWAS signals among the three populations. Firstly, the Z-score, described previously by Ioannidis et al, (2001), and by Cappelleri et al, (1996), measures the difference in estimated log-odds-ratios between the two populations divided by the estimated standard error of the difference. A ‘Z-score’ flag was set ‘on’ (value=1) if the Z-score was nominally significant at the 5% level (abs(Z)>1.96), and set ‘off’ otherwise (value=0). Secondly, an ‘opposite direction’ flag was set ‘on’ (value=1) if the odds ratios deviated from 1 in different directions (value=0 otherwise). Finally, a ‘two-fold difference in same direction’ flag was set ‘on’ (value=1) if the odds ratios were in the same direction but differed by more than two-fold between the two populations.

We assessed trans-ethic GWAS signal differences under the following “target SNP” scenarios:

1. “Causal SNP scenario”: the causal disease SNP is assumed to be known, and so is assessed directly (mimicking replication studies following a highly-powered European GWAS);
2. “Lead SNP scenario”: The causal disease SNP is assumed not known, but the genotypes of all common SNPs are assumed known (mimicking a high-density genotyping panel, perhaps together with accurate imputation). The European lead SNP (with the lowest p-value in EUR) is assessed as the target SNP (mimicking a situation where the European GWAS was performed first and the European replication study contributed to identification of the lead SNP, perhaps through combined meta-analysis and fine-mapping);
3. “Lead SNP in Illumina array scenario”: Imputation is not performed (mimicking an earlier GWAS study, or a study for which imputation is deemed unreliable). The European lead SNP is defined based on SNPs present on a representative medium-coverage GWAS panel (here, the Illumina Human Omni Express Bead Chip array), and this SNP is taken forwards for assessment as the target SNP.

## Results

A total of one thousand simulation runs, generating case-control genotype data in populations of European, Asian and Sub-Saharan African origin, were performed in this study. Logistic associations and trans-ethnic replications of GWAS signal comparisons were carried out on eligible simulated datasets as described below.

### Eligible simulations and allele frequencies

Simulations containing monomorphic target SNPs in at least one population (Asians or Sub-Saharan Africans) were not considered eligible. The number of eligible simulations varied according to the target SNP scenario considered (Table 1). Using *causal SNP*s, one hundred and forty-two simulations were eliminated due to the presence of monomorphic markers; as a consequence, 858 simulations were considered in the subsequent analyses. The allele frequencies of the simulated causal SNP across the 858 simulations showed a similar average frequency in the three populations groups (Europeans=0.35±0.24; Asians=0.36±0.28; Africans=0.35±0.27). Using *lead SNPs* with the lowest p-value in Europe as the target SNP, 807 simulations were considered eligible. The average frequency again showed similar values in the three populations groups (Europeans=0.39±0.23; Asians=0.39±0.28; Africans=0.38±0.28). Finally, considering *lead SNPs included in the Human Omni Express Bead Chip array*, the eligible simulations were a total of 797, again with similar average frequency in the three populations analysed (Europeans=0.39±0.24; Asians=0.41±0.28; Africans=0.38±0.27). Logistic association results and allele frequencies are included in Supplementary Tables 1-3.

**Table 1.** Eligible simulations and mean target SNP allele frequencies.

| Target SNP scenario | Eligible simulations / mean allele frequency (± SD) |  |  |  |
| --- | --- | --- | --- | --- |
|  | N | EUR | ASN | AFR |
| **Causal SNPs** | 858 | 0.35 ± 0.24 | 0.36 ± 0.28 | 0.35 ± 0.27 |
| **Lead SNPs** | 809 | 0.39 ± 0.23 | 0.40 ± 0.28 | 0.38 ± 0.28 |
| **Lead SNPs in Illumina array** | 797 | 0.39 ± 0.24 | 0.41 ± 0.28 | 0.38 ± 0.27 |

### Trans-ethnic differences in significant GWAS signals

Table 2 summarises the rate at which notable trans-ethnic differences were generated in our simulation study, compared to the rates seen in the study by Ntzani *et al.* (2012). The ‘Z-score’ and ‘opposite direction’ rates are expressed as a percentage of all non-null comparisons. The ‘two-fold difference’ rates are expressed as a percentage of all same-direction comparisons. Considering *causal SNPs* as the target SNP, between-population differences in estimated effect size (as measured by a nominally significant Z-score) were seen in 5.24%, 4.90%, and 5.94% of the European-Asian, European-African, and Asian-African comparisons respectively. *Lead SNPs* showed significant differences in 17.10%, 20.20%, and 10.78% of the European-Asian, European-African, and Asian-African comparisons respectively. Considering *lead SNPs included in the Illumina array*, effect size estimates differed significantly in 30.24%, 34.00%, and 8.41% of the above comparisons, which reflected a rate approximately six to seven times greater than in the *causal SNP* scenario using the European simulations.

**Table 2.** Percentage of simulations showing notable differences in effect size and/or direction of effect across populations and Ntzani et al. results.

| Reference SNPs | EUR-ASN |  |  | EUR-AFR |  |  | ASN-AFR |  |  |
| --- | --- | --- | --- | --- | --- | --- | --- | --- | --- |
|  | Z-scores | Opp. Dir. | 2-fold diff. | Z-scores | Opp. Dir. | 2-fold diff. | Z-scores | Opp. Dir. | 2-fold diff. |
| Causal SNPs | 5,24 | 1,98 | 1,07 | 4,90 | 1,17 | 0,00 | 5,94 | 3,15 | 0,96 |
| Lead SNPs | 17,10 | 8,92 | 0,95 | 20,20 | 9,67 | 0,14 | 10,78 | 11,52 | 0,56 |
| Lead SNPs in the Illumina array | 30,24 | 30,99 | 1,09 | 34,00 | 30,24 | 0,36 | 8,41 | 34,50 | 0,77 |
| Ntzani results | 21,65 | 17,53 | 38,75 | 41,67 | 20,83 | 57,89 | 23,08 | 38,46 | 50,00 |

The overall rate at which ‘opposite direction’ signals were seen was less than for ‘Z-score’ signals, but showed a similar trend towards higher frequency in the scenarios including tag-SNP effects. In the *causal SNP* scenario, the rates in the European versus Asian, European versus African and Asian versus African comparisons were 1.98%, 1.17%, and 3.15% respectively. Considering *lead SNPs*, the rates were 8.92%, 9.67%, and 11.52% respectively. Finally, the rates in the *lead SNPs included in the Illumina panel* scenario were 30.99%, 30.24%, and 34.50% respectively.

The overall rate at which ‘two-fold difference’ signals were seen was the smallest of all trans-ethnic signal difference measures, and did not show a trend towards higher frequency in the scenarios including tag-SNP effects. In the *causal SNP* scenario, the rates in the European versus Asian, European versus African and Asian versus African comparisons were 1.07%, 0.00%, and 0.96% respectively. In the *lead SNP* scenario, the rates were 0.95%, 0.14%, and 0.56% respectively. In the *lead SNPs included in the Illumina panel* scenario, the rates were 1.09%, 0.36%, and 0.77% respectively.

### Comparison to Ntzani et al. study

Ntzani and colleagues (Ntzani et al. 2012) used the NHGRI-EBI catalog of published GWAS hits (https://www.ebi.ac.uk/gwas/home) to characterize the frequency and magnitude of between-population differences in replications of GWAS signals. A total of 97 associations were evaluated in both European and Asian populations, 24 in both European and African populations, and 13 in all three groups. The percentages of these signals displaying evidence for between-population differences are displayed in Table 2.

The rate at which trans-ethnic differences were found in the Ntzani *et al.* (2012) study is considerably higher than what we found in our simulation study under the *causal SNP* and *lead SNP* scenarios. The results for the *lead SNPs in the Illumina array* scenario are more complex, with patterns varying according to which measure of trans-ethnic difference is considered, and which two populations are compared.

## Discussion

The current study was designed to allow a direct comparison to the study of Ntzani *et al.* (2012), which investigated between-population differences in replication of GWAS signals in real data collated from the NHGRI-EBI GWAS Catalogue. There are some striking differences in the rate at which between-population differences were generated in our simulation study and the generally much higher rates observed in the Ntzani *et al.* (2012) study. The rates of notable differences detected in our simulations are highly dependent on the scenario considered, thus each scenario needs to be considered separately.

The *casual SNP* scenario shows an overall consistency in the direction of effect across ancestries. This scenario generates very little between-population differences. Indeed, the rate of Z-score hits is in line with that expected under the null (5%), suggesting that between-population allele frequency differences at the true causal SNP itself are insufficient on their own to generate unusual between-population differences in estimated effect sizes.

The *lead SNP* scenario results in a noticeable increase in between-population differences. We propose that these differences are the result of (a) the Euro-centric SNP selection process and (b) between-population differences in the LD patterns between the causal SNP and the lead SNP. However, the simulated rate of between-population differences still remains considerably lower than that observed in the Ntzani *et al.* (2012) study.

However, the *lead SNPs in Illumina array* scenario leads to results similar to the Ntzani *et al.* (2012) study. We therefore propose that a combination of Euro-centric SNP selection bias with a low-coverage GWAS platform (which accentuates the between-population LD differences) is sufficient to generate rates of Z-score ‘hits’ and opposite-direction ‘hits’ without the need to posit genuine between-population differences in casual effect sizes.

We note that the extremely high rate of two-fold differences reported in the Ntzani *et al.* (2012) study requires some other explanation. We also note that the rate of significant Z-score differences seen in the Ntzani *et al.* (2012) study is considerably less than the rate of two-fold differences, which can only mean that the sample sizes used in the replication studies reported in the Ntzani *et al.* (2012) study were much lower than those used in our simulations (1000 cases and 1000 controls). For this reason, we would argue that the absolute rate of two-fold differences should be disregarded in this comparison, as presumably we could obtain the same rate simply by lowering the number of cases and controls used in our study.

In summary, our simulation study suggests that between-population differences in real causal effect size are not needed to explain the results seen in the Ntzani *et al.* (2012) study. Rather, a combination of older, low density GWAS panels, together with a strong Euro-centric bias in proposing the lead SNP, is a reasonable explanation for these results. We therefore make two predictions based on our study. Firstly, we predict that the observed between-population differences in effect size will decrease as bigger GWAS studies, with more dense panels and with better imputation, are applied. Secondly, we predict that trans-ethnic mapping will prove to be a viable method for fine-mapping, since on the one hand we propose that most causal effect sizes are shared between populations, while on the other hand we find that LD differences between populations made a considerable impact on tag-SNP effect sizes.

## Supplementary Material

**Supplementary Tables 1-3**. Simulation results for all eligible simulations. **Supplementary Table 1**: results for *causal SNP* scenario. **Supplementary Table 2**: results for *lead SNP* scenario (lowest p-value in EUR with MAF>1%). **Supplementary Table 3**: results for *lead SNP in Illumina array* scenario (lowest p-value in EUR present in the Human OmniExpress Bead Chip array). See “Headings” tab for details.

